# Age-related alterations in functional connectivity along the longitudinal axis of the hippocampus and its subfields

**DOI:** 10.1101/577361

**Authors:** Shauna M. Stark, Amy Frithsen, Craig E.L. Stark

**Affiliations:** Department of Neurobiology and Behavior, University of California Irvine

**Keywords:** hippocampus, functional connectivity, aging, anterior, posterior

## Abstract

Aging causes hippocampal circuit alterations that differentially affect hippocampal subfields and are associated with age-related memory decline. Additionally, functional organization along the longitudinal axis of the hippocampus has revealed distinctions between anterior and posterior (A-P) connectivity. Here, we examined the functional connectivity (FC) differences between young and older adults at high-resolution within the medial temporal lobe network (entorhinal, perirhinal, and parahippocampal cortices), allowing us to explore how hippocampal subfield connectivity across the longitudinal axis of the hippocampus changes with age. Overall, we found reliably greater connectivity for younger adults than older adults between the hippocampus and PHC and PRC. This drop in functional connectivity was more pronounced in the anterior regions of the hippocampus than the posterior ones, consistent for each of the hippocampal subfields. Further, intra-hippocampal connectivity also reflected an age-related decrease in functional connectivity within the anterior hippocampus in older adults that was offset by an increase in posterior hippocampal functional connectivity. Interestingly, the anterior-posterior shift in older adults between hippocampus and PHC was predictive of lure discrimination performance on the MST, suggesting that this shift may reflect a compensation mechanism that preserves memory performance. While age-related dysfunction within the hippocampal subfields has been well-documented, these results suggest that the age-related A-P shift in hippocampal connectivity may also contribute significantly to memory decline in older adults.

## 1. Introduction

The hippocampus and surrounding medial temporal lobe regions are critically involved in episodic memory (1), with rich anatomical projections to other cortical and subcortical structures that serve to guide memory behavior (2). The hippocampus itself is comprised of a network of subregions, including CA1, CA3, dentate gyrus (DG), and subiculum that are organized in a functional circuit. Projections into the hippocampus from the medial temporal lobe fall into two parallel pathways: the perirhinal cortex (PRC), which receives input from cortical areas concerned with object identity (often called the “what” pathway) and the parahippocampal cortex (PHC), which receives input from cortical areas concerned with the spatial context (often called the “where” pathway) (3, 4). These pathways converge on the entorhinal cortex (ERC), which then relays it to the hippocampus via two alternative pathways: the monosynaptic pathway, connecting the ERC and CA1 directly, and the trisynaptic pathway, with projections from ERC to DG (via the perforant path) into CA3 and then into CA1 and subiculum (5). The subiculum, along with CA1, then provides the dominant outflow from the hippocampus through deep layers of the entorhinal cortex and then the parahippocampal gyrus (6, 7).

This anatomical connectivity has important implications for memory function, such as pattern separation processes resulting from the sparse coding in the dentate gyrus and pattern completion processes derived from the recurrent collaterals in the CA3 (8, 9). However, functional organization along the longitudinal axis of the hippocampus has been a growing interest in recent years (10), which may interact with both the hippocampal subfields and their external connectivity. Organization along the long axis of the hippocampus has been demonstrated in studies of gene expression (11–13), functional neuroimaging in humans (14), and electrophysiological studies (15–17) and place field activity (18, 19). Projections from CA1 to subiculum are organized along both the transverse axis and the septo-temporal axis of the hippocampus (20), while the DG and CA3 are connected along the entire extent of the transverse axis (21). The projection from CA3 to CA1 includes a dispersion of connections in the septo-temporal axis through the Shaffer collaterals (22). This anatomical connectivity structure results in two distinct patterns: CA3 and DG tend to intermix information, whereas the CA1-subiculum projection segregates information (23).

Outsides of the hippocampus, the input from the entorhinal cortex is also organized in an anterior-to-posterior (A-P) gradient, which is largely preserved in its connections to the hippocampus (24). For example, the amygdala projects to the anterior entorhinal cortex, which then connects with the anterior hippocampus, while visual information is funneled through perirhinal and parahippocampal cortices into the posterior entorhinal cortex and then to the posterior aspects of the hippocampus. Based on these observations, the theory has been proposed that the posterior extent of the hippocampus is more likely to be involved in memory retrieval, spatial memory and navigation, while the anterior extent may be more involved in stress, emotion, goal-directed activity, and memory encoding (11, 12, 25–29). Moreover, the anterior-posterior axis of the hippocampus may interact with two larger cortical systems to support memory-guided behavior: an anterior temporal system that includes the PRC, lateral orbitofrontal cortex, amygdala, and ventral temporopolar cortex, and a posterior medial system that includes the PHC, retrosplenial cortex, and other “default network” regions (2).

Functional connectivity analyses of functional magnetic resonance imaging (fMRI) provides a popular method to evaluate correlated brain activity across structures (30, 31). In functional connectivity analyses, correlations of activity across regions over time results in maps of intrinsic functional connectivity, both across vast cortical regions (32) and within the medial temporal lobe memory system (33, 34). Previously, across 10 independent datasets, we found a striking pattern of sub-networks within the MTL: 1) high connectivity between the hippocampal subfields bilaterally, 2) high connectivity between entorhinal-perirhinal cortices bilaterally, and 3) high connectivity within the parahippocampal cortex bilaterally (33). We were struck by the difference between the functional connectivity (consistent across task-based and resting state fMRI) and the connectivity predicted by the anatomical structure. While these data emphasize connectivity within the hippocampal circuit between subfields, they do not address potential interactions with segregation along the longitudinal axis of the hippocampus. However, Libby et al. (35) showed greater connectivity between PRC and the anterior hippocampus, whereas there was greater connectivity between PHC and posterior hippocampus in young adults. As for the subfields, CA1 and subiculum mirrored this A-P pattern with PRC and PHC, but this connectivity gradient with DG/CA3 was less robust.

Hippocampal subfields undergo age-related changes in specific ways that impact memory decline in older age and may also impact functional connectivity (36). Age-related volumetric declines have been observed, cross-sectionally, in the DG/CA3 and CA1 (37–39). Age-related changes within this hippocampal circuit may significantly contribute to memory decline and connectivity to other cortical regions (40). Specifically, the dentate gyrus receives reduced inputs from the entorhinal cortex (41) and is coupled with a decrease in modulation by inhibitory neurons (42) leading to a hypoactive DG. In contrast, a decrease in cholinergic modulation in normal aging (43) creates hyperactivity in CA3, which has been observed in high firing rates in animals (44) and fMRI signal in humans (45, 46). Finally, dopaminergic projections to CA1 are reduced in aging (47), which may contribute to a shift in the balance of projections from CA3 to the entorhinal cortex leading into CA1.

It is unknown how these age-related changes to the hippocampal circuit may be differentially impacted along the longitudinal axis of the hippocampus. A whole-brain analysis contrasting young and older adults (48) found diminished functional connectivity to neocortical areas in older adults, but a relative increase in posterior hippocampal-neocortical connectivity in aging. These findings are consistent with observations used to support the Posterior-Anterior Shift in Aging (PASA) model (49), where whole brain functional activity is decreased in anterior regions relative to posterior regions (50), possibly reflecting compensation mechanisms that predict preserved memory performance. However, it is unclear if this anterior-to posterior (A-P) shift in aging is systematic across the hippocampal subfields or if DG/CA3 is relatively preserved given previous findings reflecting the more integrated nature of its connectivity (35).

Here, we sought to examine the functional connectivity differences between young and older adults at high resolution within the MTL network, allowing us to explore how hippocampal subfield connectivity across the longitudinal axis of the hippocampus changes with age. We assessed functional connectivity primarily during an incidental encoding task (two other task-related datasets are included for comparison), providing an opportunity for hippocampal engagement in both young and older adults. Resting state scans suffer from the potential confound that young and older adults may typically engage in different tasks or thought patterns while resting, potentially introducing enough variance to obscure true differences and clouding interpretation of any differences that do come through. Consistent with this observation, in our previous work, we did not observe an age-related difference in the functional connectivity matrix for resting state data, but we did when considering functional connectivity from task-related activity (33). By constraining task demands, we can eliminate this potential source of variability, which may significantly contribute to age-related differences in the default mode network (51).

Specifically, we first sought to evaluate functional connectivity along the longitudinal axis of the hippocampus to the ERC, PRC, and PHC in both young and older adults. We anticipated replicating earlier findings, showing greater connectivity in older adults between anterior hippocampus and PHC (48). In addition, we predicted age-related declines in functional connectivity in anterior regions compared to posterior ones, consistent with previous findings of an A-P shift in older adults. Next, we evaluated hippocampal subfield connectivity to the MTL network in both young and older adults, hypothesizing reduced connectivity of DG/CA3 in older adults given age-related functional activity dysfunction in this region (52). Finally, we explored functional connectivity along the longitudinal axis of the hippocampus for each of the subfields, taking into account the disproportionate representation of these regions along the A-P axis.

## 2. Materials & Methods

### 2.1. Participants

A total of 31 young (18F/13M; mean age = 29 years; range = 20-39) and 31 older (16F/15M; mean age = 76 years; range = 70-86) adults were recruited from the University of California, Irvine and surrounding area. They provided written consent in compliance with the UCI Institutional Review Board and were compensated for their participation. All participants were screened for history of neurological disease or psychiatric illness, spoke fluent English, were right-handed, had normal or corrected-to-normal vision, and no contraindications for MRI. To determine cognitive status, all participants also completed the Mini Mental State Exam (53). Young participants scored 29-30 and older adults scored 27 or above, all in the normal range for their age.

### 2.2. Mnemonic Discrimination Task (MST)

During a behavioral testing session separate from the scan date, participants completed the MST (publicly available here: https://faculty.sites.uci.edu/starklab/mnemonic-similarity-task-mst/) using sets C & D. In the first phase, participants engaged in an incidental encoding task consisting of an indoor/outdoor judgment for each object (based on their opinion with no right or wrong answer) via a button press (128 items total, 2s object presentation and 3s scene presentation time, 0.5s ISI). Immediately following the encoding task, participants were shown video instructions describing the test phase to ensure consistent delivery of the instructions. In the test phase, they engaged in a modified recognition memory test in which they identified each item as “Old”, “Similar”, or “New” via button press (192 items total – 64 repeated items, 64 lure items, and 64 foil items; 2.5s/3s each, ≥0.5s ISI). The image disappeared from the screen after 2.5s (or 3s in the case of scenes), replaced by a white screen until participants responded. This enables participants to respond at whatever pace they feel comfortable with but does not allow for variable time in inspecting the image. One-third of the images in the test phase were exact repetitions of images presented in the encoding phase (targets or repeats); one-third of the images were new images not previously seen (foils); and one-third of the images were similar to those seen during the encoding phase, but not identical (lures). These trial types were randomly intermixed during the test.

As in our prior work (Stark et al., 2013; 2015; 2017), the Lure Discrimination Index (LDI) was calculated as the difference between the rate of “Similar” responses given to the lure items minus “Similar” responses given to the foils (to correct for any response biases). Recognition (REC) for repeat items was calculated as the difference between the rate of “Old” responses given to repeat items minus “Old” responses given to foils (aka “corrected recognition memory scores”). These scores correct for any response bias on a per-subject basis.

### 2.3. fMRI task

Each of 8 fMRI runs consisted of 96 objects presented in a continuous incidental encoding paradigm that mirrors the MST but does not have an explicit memory demand. This task is virtually identical to the task we have used several times previously (54, 55). These items were different from those viewed in the behavioral MST. Each item was presented for 2.4 seconds, followed by an inter-trial interval of 0.29 seconds. Participants were instructed to indicate if each item was an “indoor” or an “outdoor” item, with an emphasis that there was no correct answer and to simply respond with their best guess. 384 of these items were novel foils that were never repeated while 96 items were later repeated (repeats) and 96 additional items were repeated with items that were similar, but not exactly the same (lures). In total, there were 768 trials divided into 8 runs of 96 trials each.

### 2.4. Imaging Parameters

All scanning was performed on a Phillips 3.0 Tesla Scanner (Best, the Netherlands), using a 32-channel sensitivity encoding (SENSE) head coil at the Research Imaging Center at UC Irvine. During each of 8 scanning runs, 173 T_2_*-weighted, single-shot echo-planar volumes were acquired angled parallel to the long axis of the hippocampus and covering most of the MTL in 22 slices. Each slice was 1.3 mm thick separated by a 0.2 mm gap. Functional pulse sequences had a repetition time (TR) of 1500 ms, an echo time (TE) of 26 ms, a flip angle of 70°, an acquisition matrix size of 128 × 128 mm, a field of view (FOV) of 180 × 180 mm, and a SENSE factor of 2.5, resulting in an in-plane resolution of 1.5 × 1.5 mm (scan resolution is 120×120). The first four functional volumes were discarded to accommodate for T1 equalization.

Additionally, T1-weighted whole-brain anatomical images were acquired using a sagittal magnetization-prepared rapid gradient echo (MP-RAGE) scan (TR 11ms; TE 4.6 ms; flip angle 18°; matrix size 320 × 320 mm; FOV 240 × 240 × 150 mm; resolution 0.75 mm isotropic; 200 slices).

### 2.5. fMRI Pre-processing

We used Analysis of Functional Neuroimages (AFNI; version 17.1.09)(56) software to perform most of the imaging data analyses. Functional data for these analyses were slice-time and motion corrected using six rigid-body transformation parameters using the function align_epi_anat.py (57). In order to reduce the effects of physiological noise in the BOLD signal, we passed the filtered data through ANATICOR (58) using the white matter and CSF maps created from Freesurfer. We used AFNI’s 3dTproject function to apply a temporal bandpass filter of 0.009 to .08, to censor TRs that had excessive motion (0.5mm and/or 0.5 degrees) and the following TR, and to regress out the global mean signal for each run. While there is considerable debate regarding the removal of the global signal due (59–61), we reasoned that removing the global mean signal would better address spurious and inflated correlations due to motion. Further, we are focusing here on relative differences in connectivity between groups rather than the degree of connectivity per se, which should be preserved here even if negative correlations were induced by the removal of the global mean signal. Each subject’s anatomical image was segmented into grey matter, white matter, and cerebral spinal fluid probability (CSF) maps using Freesurfer (62, 63). Each run was then concatenated into a single time series for each participant.

Both the whole-brain and high-resolution functional data were aligned to the participant’s MP-RAGE using the script align_epi_anat.py (57) from AFNI (56). Each participant’s structural scan and functional data (statistical maps) were aligned to a model template using ANTs (Advanced Normalization Tools; (64), which normalizes each participant’s T1-weighted MP-RAGE to a template space based on MNI coordinates but derived from previous work in our lab (55). Briefly, it is based on a central tendency of 20 healthy adults (including younger and older adults) previously aligned to MNI space. ANTS combines a 12-parameter affine registration with a diffeomorphic 3D vector field mapping (Syn) to perform invertible, smooth mappings between the original participant space and the model template space.

To segment the MTL and calculate MTL volumes for individuals in the current study, we created a multi-atlas model using ASHS (65) and 19 independent hand-segmented brains (both the MP-RAGE and high-resolution T2 images). These segmentations included both segmentations of the parahippocampal gyrus into perirhinal (PRC), parahippocampal (PHC), and entorhinal (ERC) cortices described previously (66, 67). Similarly, we segmented the hippocampus into 3 regions: a combined dentate gyrus and CA3 (DG/CA3), CA1, and subiculum, based on our previous work (Stark & Stark, 2017). For each of these, we created multi-atlas models in ASHS and then used these to segment each individual included here. Briefly, ASHS performs a non-linear registration between a new participant’s structural scan and each of the scans in the multi-subject atlas using ANTS (64). A voting procedure was then enacted to determine an initial segmentation based on the degree of deformation needed to warp the new participant onto each atlas (less deformation needed implies a higher degree of match and therefore a greater weight of that individual atlas to the segmentation). Finally, an AdaBoost technique was used to detect and remove segmentation biases.

To examine the long-axis, we split the hippocampal segmentations into six regions (Figure 1A). First, we divided the template-space brain into equally spaced slices along a roughly 30 degrees forward-facing angle (aligned with the angle of the hippocampus), resulting in 6 segments with roughly the same number of voxels, separated by small gaps to reduce partial volume effects. Then we multiplied this template into subject-specific space creating even segmentations along the longitudinal axis of the hippocampus.

**Figure 1.**
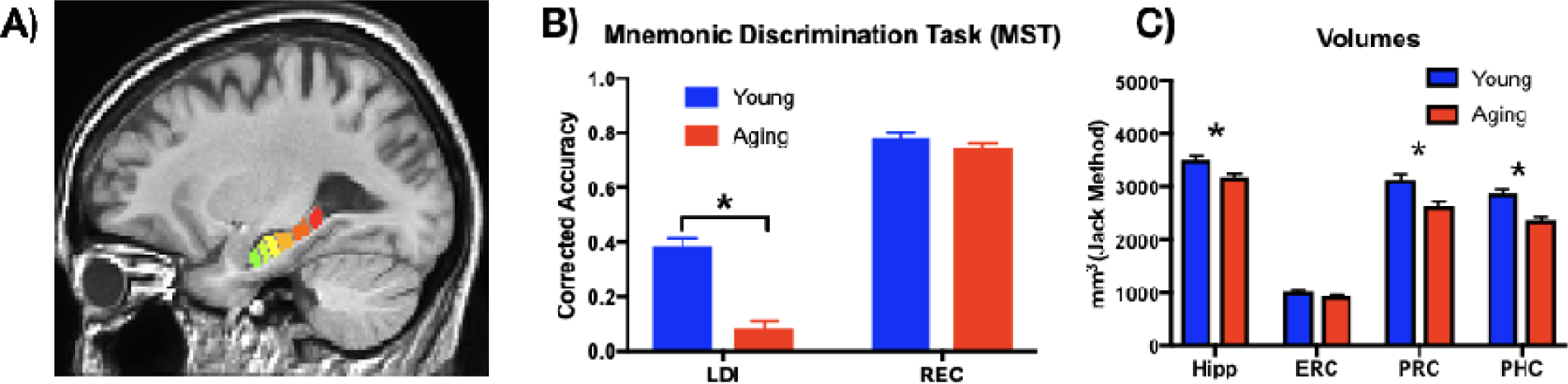
A) Segmentation of the hippocampus into 6 roughly equivalent regions along the anterior-posterior axis. B) Behavioral performance on the MST demonstrating impaired lure discrimination performance in older adults, but intact recognition performance. LDI – lure discrimination index; REC – recognition memory index C) Younger adults had greater volume than older adults in the hippocampus, PRC, and PHC. Hipp – hippocampus, ERC – entorhinal cortex, PRC – perirhinal cortex, PHC – parahippocampal cortex. * p < .05

Finally, intracranial volume was calculated by a Freesurfer (68) segmentation of the MP-RAGE based on a combination of white matter, gray matter, and cerebral spinal fluid. To adjust for head size and age, we calculated an adjusted volume for each structure in cubic millimeters (mm^3^), such that the adjusted Volume = raw volume - b x (ICV - mean ICV), where b is the slope of the regression of an ROI volume on ICV (69). We calculated the slope of the regression for each ROI and the mean ICV based on a larger lifespan dataset (n= ∼160, ages 20-89 years old) that we have published on previously (70, 71). The adjusted volumes were calculated for the left and right ROIs separately and then averaged.

### 2.6. Functional Connectivity Analyses

To assess functional connectivity between the hippocampus and medial temporal lobe, we employed ROI-to-ROI correlation analyses. Functional connectivity refers to the temporal correlations of the time series between spatially separated brain regions. Using 3dTproject (AFNI), we extracted the average residual time-series across the task for each ROI. Pearson’s r correlation coefficients were then calculated between regions and then Fisher’s r-to-z transformed. All further statistical analyses were performed using Prism 7.0d (www.graphpad.com).

## 3. Results

### 3.1. MST behavioral results

We tested for a replication of our earlier findings in each measure from the MST independently: namely, matched recognition performance between older and younger adults, with an age-related decrease in LDI performance. Therefore, LDI and REC scores for younger and older adults were entered into a 2×2 ANOVA (Prism 7.0d), with factors of Age and Test Type (Figure 1B). We observed a main effect of Age (F(1,60) = 52.1, p<.0001), a main effect of Test Type (F(1,60) = 462.1, p<.0001), and an interaction (F(1,60) = 29.38, p<.0001). Bonferroni-corrected multiple comparisons showed no difference in REC (Young=0.78 vs Aging=0.74), but a significant reduction in LDI for older (0.08) compared to younger (0.38) adults (t(120) = 8.9, p<.0001). Thus, consistent with our previous findings (39, 72, 73), older adults reliably demonstrate greatly reduced lure discrimination performance for related lure items, while recognition memory for repeated items remained intact.

### 3.2. Functional connectivity along the longitudinal axis of the hippocampus

To evaluate whether connectivity along the long axis of the hippocampus differed for young and older adults, we calculated functional connectivity along the 6 hippocampal (Hx) segmentations to three separate medial temporal lobe regions, averaging across left and right hemispheres: ERC, PRC, and PHC. These data were entered into a 2×6 repeated-measures ANOVA with age (young or aging) and region (1,2,3,4,5,6) as variables for each region. All three regions (ERC, PRC, and PHC) showed a main effect across the six hippocampal subdivisions (main effect of region: ERC: F(5,300) = 3.1, p<.01; PRC: F(5,300) = 6.8, p<.0001; PHC: F(5,300) = 7.8, p<.0001). While ERC showed no main effect of age or interaction (Supplemental Figure 1D), we observed (Figure 2A-B) greater connectivity between the Hx and PHC (F(1,60) = 11.1, p <.01) for younger than older adults and only modest evidence for a similar effect of age in the PRC (F(1,60) = 3.4, p = .07). In addition, FC with PHC changed across the six hippocampal subdivisions with a significant interaction (F(5, 300) = 3.0, p<.014). Šidák’s multiple comparisons tests on each of the 6 regions revealed greater FC in young than older adults for the three most anterior regions (1: t(360) = 2.9, p<.02; 2: t(360) = 4.7, p<.0001; 3: (t(360) = 2.7, p <.05), but not the three most posterior regions (4: t(360) = 2.3, p=.13; 5: t(360) = 1.9, p=.31; 6: t(360) = 1.5, p=.54).

**Figure 2.**
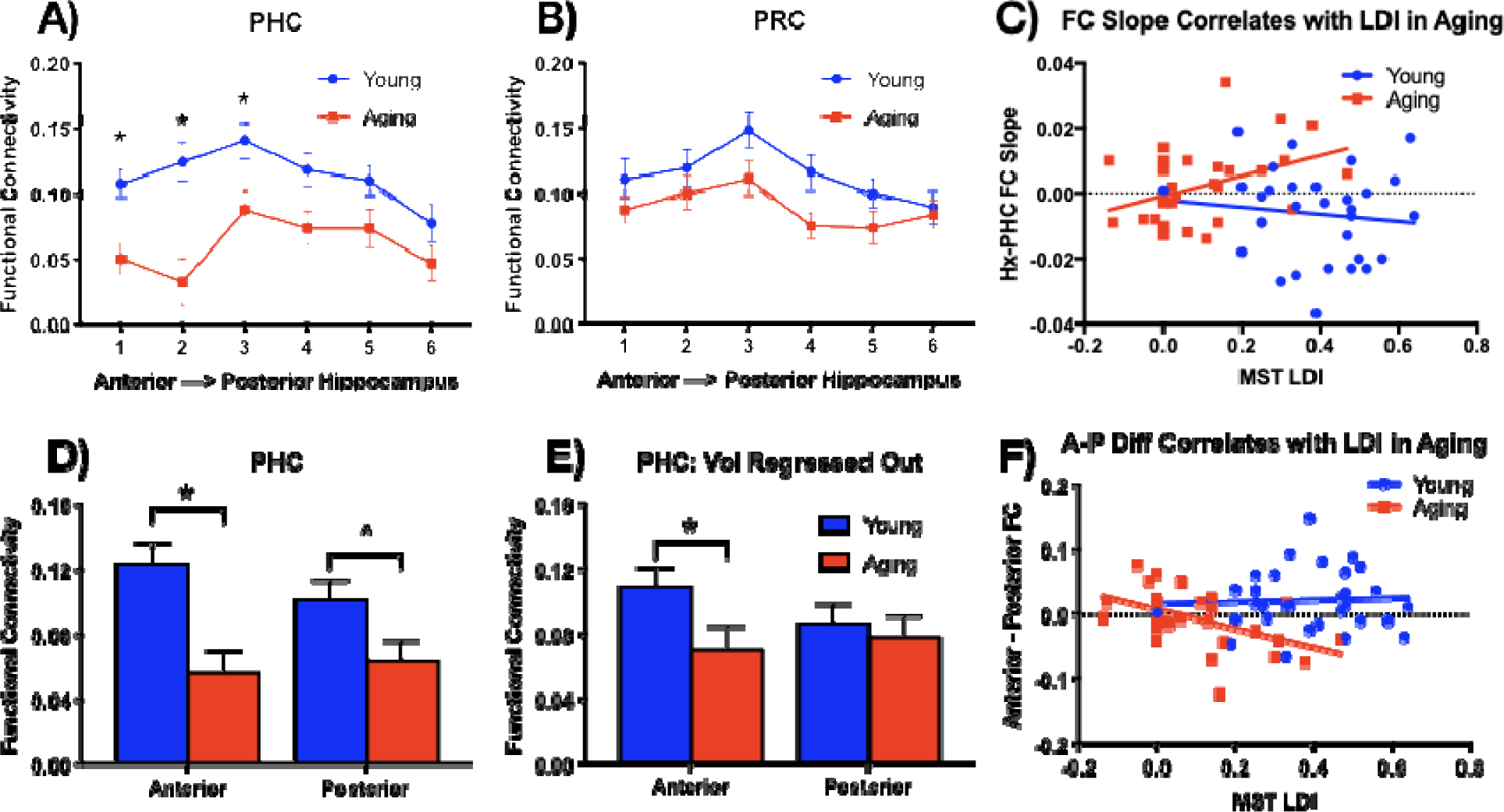
A) Functional connectivity between the hippocampus and PHC is greater for young than aging adults in the anterior portions of the hippocampus. B) Similar age-related decrease in FC between Hx-PRC, but no difference across the longitudinal axis of the hippocampus. C) The Hx-PHC FC connectivity slope along the longitudinal axis of the hippocampus predicts lure discrimination performance (LDI) in older adults, but not in younger adults. D) Focusing on the anterior (first 3 subregions) and posterior (last 3 subregions) again emphasized the interaction, such that there is a greater age-related drop in FC for older adults in the anterior regions of the hippocampus. E) Regressing out volumes of the hippocampus and PHC did not change this overall effect, eliminating concerns over reduced FC due to volumetric declines. F) A greater shift from anterior to posterior FC predicted greater lure discrimination performance in older adults, but not in younger adults. * p < .05, ^ p = .06

Since we do not have strong predictions about each of these 6 regions independently, to more directly explore the anterior vs. posterior hippocampal dissociation, we combined the hippocampal segmentations into anterior (first 3 segmentations) and posterior (last 3 segmentations) to evaluate the age difference using Šidák’s multiple comparisons test. Older adults had reduced FC with the PHC in the anterior hippocampus compared to younger adults (t(120) = 4.0, p<.001), an effect that was only weakly observed in the posterior hippocampus (t(120) = 2.2, p=.06) (Figure 2D). This effect was reflected by a significant age x region interaction (F(1,60) = 6.8, p<.02). Thus, young adults demonstrate greater FC between the Hx and PHC than older adults, particularly in the anterior regions of the hippocampus. The corresponding analysis with the PRC showed a clear effect of anterior-posterior on hippocampal connectivity (F(1,60) = 16.7, p<.001) and some evidence for a main effect of aging (F(1,60) = 3.5, p = .07), but no evidence for any interaction (p=0.87). To determine the reliability of these findings, we applied the same analyses to two additional datasets (see Supplemental Materials): another incidental encoding task (using scenes in this case) and a continuous recognition memory test. Both tested similar age ranges and in both cases, we found stronger age-related reductions in Hx-PHC FC in the anterior regions of the hippocampus than in the posterior regions (Supplemental Figure 1). In addition, consistent with our main analysis, we did not observe strong evidence for similar effects in the PRC. Thus, we have replicated these basic findings twice. In addition, we explored whether there was hemispheric asymmetry in these findings by examining each FC relationship within hemisphere (see Supplemental Materials). We found some evidence for hemispheric asymmetry, with larger effects present in the right than left hemisphere, but both hemispheres showed a consistent age-related decrease in Hx-PHC functional connectivity.

We were concerned that age-related reductions in volume might alter the FC between regions. Indeed, when we entered the corrected volumes into a 2×4 ANOVA, we found greater volumes for the young than older adults (main effect of age: F(1,60) = 57.4, p<.0001), differences in overall regional volumes (main effect of region: F(3,180) = 691.5, p<.0001) and an interaction (F(3,180) = 6.7, p<.001). Post-hoc Šidák’s multiple comparisons revealed greater volumes for young than aged adults for the hippocampus (t(240) = 3.5, p<.01), PRC (t(240) = 5.3, p<.001), and PHC (t(240) = 5.5, p<.001) (Figure 1C). Therefore, we examined FC for Hx-PHC, regressing out volume on an individual basis for both the hippocampus and parahippocampal cortex. Even when controlling for differences in volume, we found that FC changed across the six hippocampal subdivisions (main effect of region: F(5,300) = 7.8, p<.0001) and an interaction (F(5,300) = 2.9, p<.05), largely driven by greater FC in young adults for hippocampal segment 2 (t(360) = 3.2, p<.01). Combining the hippocampal segmentations into posterior (first 3 segmentations) and anterior (last 3 segmentations), we found that older adults had reduced FC in the anterior hippocampus than younger adults (t(120) = 2.3, p<.05), but not in the posterior hippocampus (t(120) = .51, p=.85) (Figure 2E). Thus, volume decreases associated with aging are not artificially altering the observation of a larger reduction in functional connectivity between the Hx-PHC in the anterior regions of the hippocampus than the posterior regions.

We were also concerned that aging might be associated with a decrease in signal to noise, perhaps from a change in vasculature (74, 75) in the MTL that might induce our observed age-related declines. Thus, we examined connectivity with the ERC as a control region because it did not reveal an age-related decline in FC. As noted above, there was no main effect of age on Hx-ERC connectivity or interaction between age and longitudinal position. Likewise, there was no reliable correlation between age and overall Hx-ERC functional connectivity (r=-0.11, p=0.23) or between age and overall PHC-FC (r=-0.12, p=0.22). Similarly, the average FC between the hippocampus and PHC was comparable to that observed with ERC in older adults (t(30) = .04, p = .97)). Thus, reduced signal to noise is unlikely to account for the age-related decrease in FC between Hx-PHC and Hx-PRC (see also Figure 3 and Discussion).

**Figure 3.**
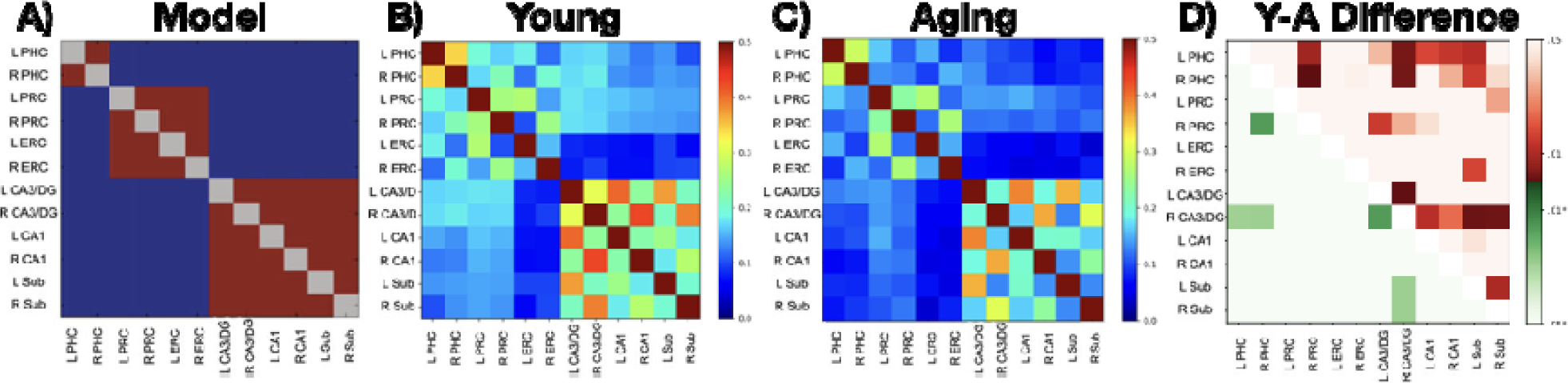
A) Model of functional connectivity based on Lacy et al. (2011). B) Functional connectivity matrix for young adults and C) older adults. D) Difference matrix testing whether regions showed greater connectivity in young than old. Values above the diagonal (reds) are uncorrected p-values while values below the diagonal (greens) reflect corrections for multiple comparisons.

Finally, we were interested in the relationship between Hx-PHC functional connectivity and lure discrimination performance on the MST. We already observed that older adults are impaired on lure discrimination relative to younger adults, which has been linked to reductions in hippocampal volume (39) and functional activity in the hippocampus (46, 52). We calculated a slope for each person across the 6 hippocampal segmentations’ connectivity with the PHC and then examined the relationship between that slope and lure discrimination performance. Older adults showed a positive relationship such that greater lure discrimination performance was predicted by a steeper slope of PHC functional connectivity along the A-P axis of the hippocampus (F(1,28) = 5.1, p<.05), while this relationship was not present in younger adults (F(1,28) = .39, p=.53) (Figure 2C). To more directly assess the age-related anterior to posterior shift in aging on behavior, we conducted a linear regression with lure discrimination on the difference in functional connectivity between the 3 anterior regions and the posterior 3 regions. We replicated the same pattern, such that the difference between connectivity in the anterior compared to the posterior hippocampus was predictive of lure discrimination ability in older adults (F(1,28) = 8.5, p<.01), but not in younger adults (F(1,28) = .06, p = .80) (Figure 2F). Thus, functional connectivity along the long axis of the hippocampus-PHC in older adults is predictive of memory performance on a hippocampal-dependent task.

### 3.3. Functional connectivity in hippocampal subfields

Connectivity along the longitudinal axis of the hippocampus does not take into account the subfield circuit known to be critical for hippocampal computations. Therefore, we sought to explore functional connectivity of the hippocampal subfields: DG/CA3 (combined due to resolution constraints), CA1, and the Subiculum with the medial temporal lobe cortices. Previously, we have shown a clear pattern of three subnetworks of functional connectivity (Figure 3A): 1) within the hippocampal subfields, 2) entorhinal-perirhinal cortices, and 3) parahippocampal cortex (33). This pattern was reliable and likely represents a pattern of intrinsic connectivity within the MTL (34), much like similar patterns observed at the whole brain level (76). Consistent with our previous findings and the model, both the Young (Figure 3B) and Aging (Figure 3C) matrices fit the same pattern: relatively high correlation coefficients amongst the hippocampal subfields, between bilateral ERC-PRC, and between bilateral PHC. Akin to our prior work, we calculated z-transformed correlation coefficients between this intrinsic connectivity model and each subject’s matrix (lower-triangle only) and between an anatomically-based model and each subject’s matrix. Both young and aged subjects correlated reliably with the prior model (young z=0.54, t(30)=15.2, p<0.0001; aged z=0.53, t(30)=16.5, p<0.0001) and not the anatomically-based model (young t(30)=1.0, p=0.31; aged t(30)=0.72, p=0.48). There was no evidence for an age-related difference in the correlation with either the intrinsic connectivity model (t(60)=0.22, p=0.83) or the anatomically-based model (t(60)=1.23, p=0.22).

Directly contrasting the young and old matrices, we identified several regions of greater FC for young adults compared to older adults by contrasting the z-transformed correlation coefficient matrices across groups using unpaired t-tests both without and with FDR-based corrections for the 66 comparisons (Figure 3D). Adjusting for multiple comparisons with an false-discovery rate (FDR) of Q = .05, younger adults had significantly greater FC in the following regions: 1) right DG/CA3 ↔ left DG/CA3 (t(31) = 3.43, p=.00055 uncorrected q=0.016), 2) right DG/CA3 ↔ left Subiculum (t(31) = 3.06, p=.0016, q=.029), 3) right DG/CA3 ↔ right Subiculum (t(31) = 2.96, p=.0022, q=.029), 4) right DG/CA3 ↔ left PHC (t(31) = 2.86, p=.0029 uncorrected, q=.029), 5) right DG/CA3 ↔ right PHC (t(31) = 2.93, p=.0024, uncorrected, q=.029), and 6) right PHC ↔ right PRC (t(31) = 3.55, p=.00038 uncorrected, q=0.016). Some evidence was also found for greater connectivity in the left PHC ↔ right PRC contrast as well (t(31) = 2.48, p=.008 uncorrected, q=0.069)). Again, we found supporting evidence for these findings in two additional datasets (Supplemental Figure 2), emphasizing an age-related decrease in functional connectivity for DG/CA3 and PHC, and to a lesser degree, CA1 and subiculum. There is also some laterality asymmetry (notably, right DG/CA3), but it was not reliable enough across datasets to draw strong conclusions. Importantly, these results demonstrate that age-related modulation of the functional connectivity can be characterized by hippocampal subfield specificity, particularly DG/CA3. We next explore how these two approaches intersect: does connectivity along the longitudinal axis of the hippocampus account for these differences in subfield connectivity?

### 3.4. Relationship between hippocampal subfields and MTL along the longitudinal axis of the hippocampus

In section 3.2, we established that there is an age-related decrease in Hx-PHC FC, particularly in the anterior hippocampus. Then, in section 3.3, we found an age-related decrease in FC between DG/CA3-PHC. Here, we explored the connectivity within each subfield (DG/CA3, CA1, and Subiculum) to PHC and PRC along the longitudinal axis of the hippocampus. We applied the 6-segmentation mask to the hippocampal segmentation mask and correlated activity in each region to the PHC and PRC, collapsing across hemispheres to increase signal-to-noise and power (Figure 4A-F). These data were entered into a 2×6 repeated-measures ANOVA with age (young or aging) and region (1-6) as variables for each subfield. For PHC, all three subfields showed greater connectivity for younger than older adults (main effect of age: DG/CA3: F(1,60) = 8.3, p <.01; CA1: F(1,60) = 7.5, p<.01; Sub: F(1,60) = 4.8, p<.05) and a main effect across the six regions for DG/CA3 (main effect of region: DG/CA3: F(5,300) = 10.10, p<.0001; Sub: F(5,300) = 4.3, p<.001). In addition, there was an interaction in CA1 (F(5, 300) = 3.5, p<.01). Again, we combined the hippocampal segmentations into anterior (first 3 segmentations) and posterior (last 3 segmentations) to evaluate the age difference using Šidák’s multiple comparisons test. In all three hippocampal subfields, younger adults had greater functional connectivity to the PHC in the anterior hippocampus than older adults (DG/CA3: t(120) = 3.3, p<.01; CA1: t(120) = 373, p<.01; Subiculum: t(120) = 2.9, p<.01). While there was a numerical decrease in the posterior regions, none of them passed the statistical threshold of p<.05.

**Figure 4.**
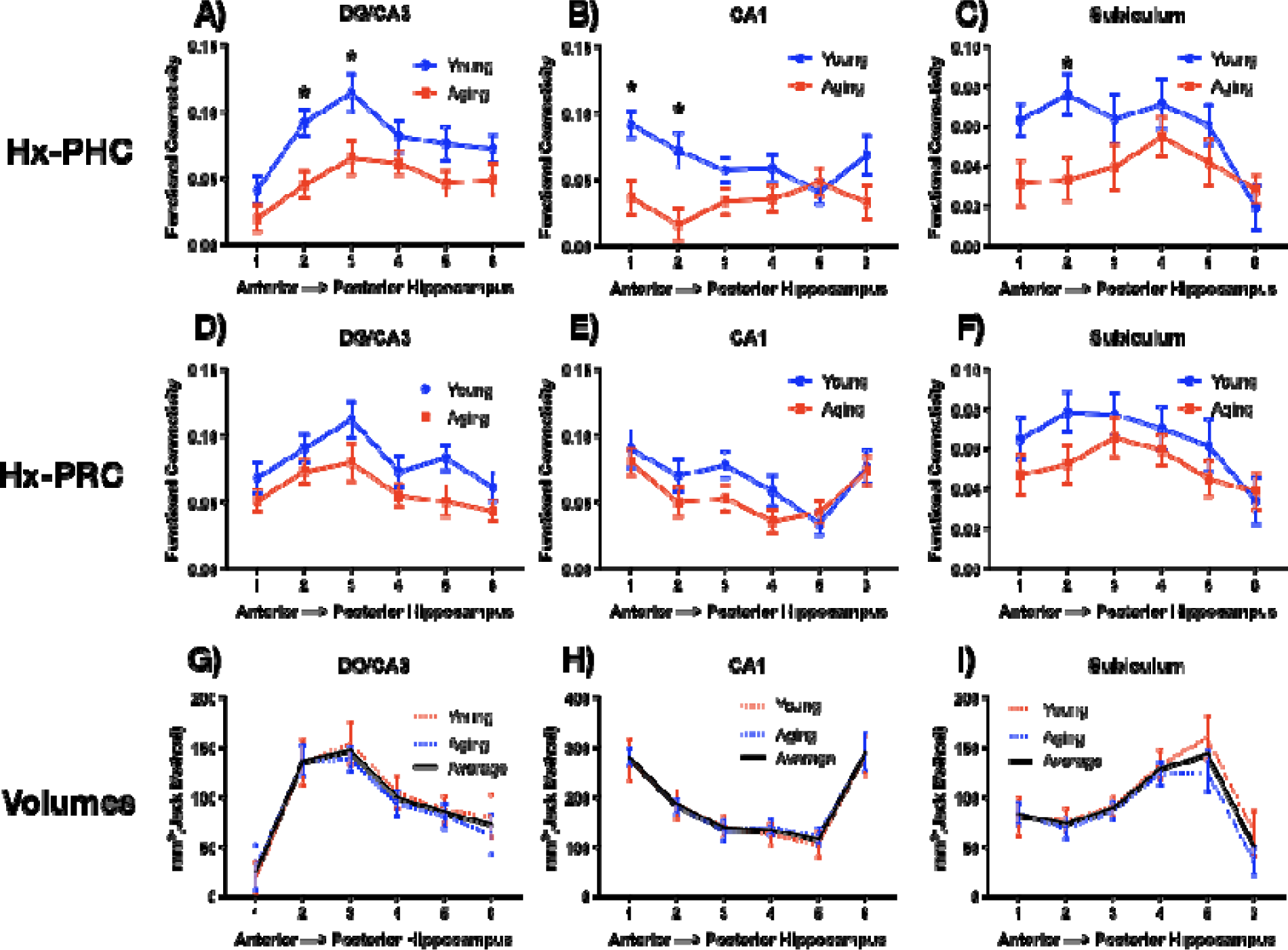
Functional connectivity was greater in young than older adults between each hippocampal subfield and PHC: DG/CA3 (A), CA1 (B), and Subiculum (C): and PRC: DG/CA3 (D), CA1 (E), and Subiculum (F). While there were fewer voxels in DG/CA3 and Subiculum in older adults, the distribution of representation across the longitudinal axis revealed more voxels in the anterior than posterior region for DG/CA3 (G), while CA1 showed a u-shaped distribution (H), and the Subiculum had a greater number of voxels in the posterior regions (I).

For PRC, only DG/CA3 showed a greater connectivity for younger than older adults (main effect of age: DG/CA3: F(1,60) = 6.9, p <.05), but all three subregions showed a main effect across the six regions (main effect of region: DG/CA3: F(5,300) = 7.0, p<.0001; CA1: F(5,300) = 7.2, p<.0001; Sub: F(5,300) = 4.4, p<.01), and no interactions. Again, we combined the hippocampal segmentations into anterior (first 3 segmentations) and posterior (last 3 segmentations) to evaluate the age difference using Šidák’s multiple comparisons test. In contrast to PHC, none of the hippocampal subfields showed greater functional connectivity in the anterior or posterior hippocampus for younger adults than older adults (Supplemental Figure 3D-F).

These subfield level differences raise the possibility that our earlier observation of functional connectivity modulation along the longitudinal axis arose not from the longitudinal axis *per se*, but from the fact that the subfields are not distributed evenly across the long axis (Figure 4G-I). Could differential changes in subfield volume with age or interactions along the long axis induce our previously-observed age-related changes in functional connectivity? For each subject, we calculated subfield volumes in each of our six long-axis segments and entered these data into a 2×6 repeated-measures ANOVA with age (young or aging) and region (1-6) as variables for each subfield. Overall, a main effect of region was found for all three subfields, demonstrating a non-uniform distributions across the long axis (DG/CA3: F(5,300) = 493.3, p<.0001; CA1: F(5,300) = 635.8, p<.0001; Subiculum: F(5,300) = 420.2, p<.0001). Younger adults had greater volume in DG/CA3 than older adults (main effect of age: F(1,60) = 6.0, p<.02), with this effect being more pronounced in the posterior hippocampus (age x region interaction F(5,300) = 7.9, p<.0001). Similar results were observed in subiculum, with an age-related reduction in volume (main effect of age: F(1,60) = 31.4, p<.0001) that was more pronounced in posterior regions (age x region interaction (F(5,300) = 18.9, p<.0001). Interestingly, there was no age-related decrease in CA1(main effect of age: F(1,60) = 1.8, p=.18).

These volumetric differences emphasize three critical points: 1) the distribution of subfields is not even across the longitudinal axis of the hippocampus, with greater representation of DG/CA3 in anterior regions and the opposite pattern in the Subiculum, while CA1 shows a more U-shaped distribution with the greatest representation at the head and tail. 2) age-related volume changes are not uniform across the hippocampus, but appear to be more specific to the DG/CA3 and subiculum, particularly in the more posterior regions of hippocampus; and 3) while age-related volume changes are greater in posterior portions of the hippocampus, our functional connectivity changes were greater in more anterior portions of the hippocampus, making this a highly unlikely account for the observed functional connectivity results. Finally, it is worth noting that the first subregion in DG/CA3 and the last subregion in the subiculum both contain very few voxels, so FC from those regions should be interpreted with caution.

### 3.5. Functional connectivity within the longitudinal axis of the hippocampus

Having established the FC profiles along the longitudinal axis of the hippocampus with the MTL, we sought to explore FC within the hippocampus itself. A recent investigation in young adults showed greater inter-voxel similarity for the anterior than the posterior hippocampus, indicating greater FC within the anterior portion (77). Similarly, we were interested in how age-related change along the longitudinal axis of the hippocampus may influence the connectivity among these regions. To mirror this prior inter-voxel similarity analysis, we first calculated the z-transformed correlation coefficients for every pair of voxels within each subject’s hippocampus and averaged these according to segment to create average segment to segment correlation scores (e.g., average of all segment 1 to segment 1 correlations, average of all segment 1 to segment 2 correlations, etc. leading to 6×6 correlation matrices). We then compared the Young and Aged groups using unpaired t-tests for the left, right, and the average of both hemispheres. (Figure 5A-C). We found a clear pattern consistent with aging modifying the pattern observed by (77). The anterior segments (1-4) exhibited greater FC for young adults than older adults (Figure 5A), particularly in the right hemisphere (Figure 5C). In contrast, we observed greater FC in the posterior regions (4-6) for older adults than young adults, particularly in the left hemisphere (Figure 5B). Similar results, though less striking, were observed in our two additional datasets, supporting anterior hippocampal dysfunction in aging that may be offset by posterior hippocampal engagement. This pattern did not appear subfield-specific as we observed a similar pattern of results across the three hippocampal subfields (CA1, DG/CA3, and Subiculum), with an age-related decrease in FC for anterior regions and an age-related increase in posterior regions (Figure 5D-F). We chose to simply set our alpha threshold at p<.05 for each comparison, recognizing that many of them would not survive corrections for multiple comparisons. However, we would be remiss to ignore these findings, particularly given the prior findings in young individuals, the exceptional regularity of this pattern across the subfields and in our additional two datasets.

**Figure 5.**
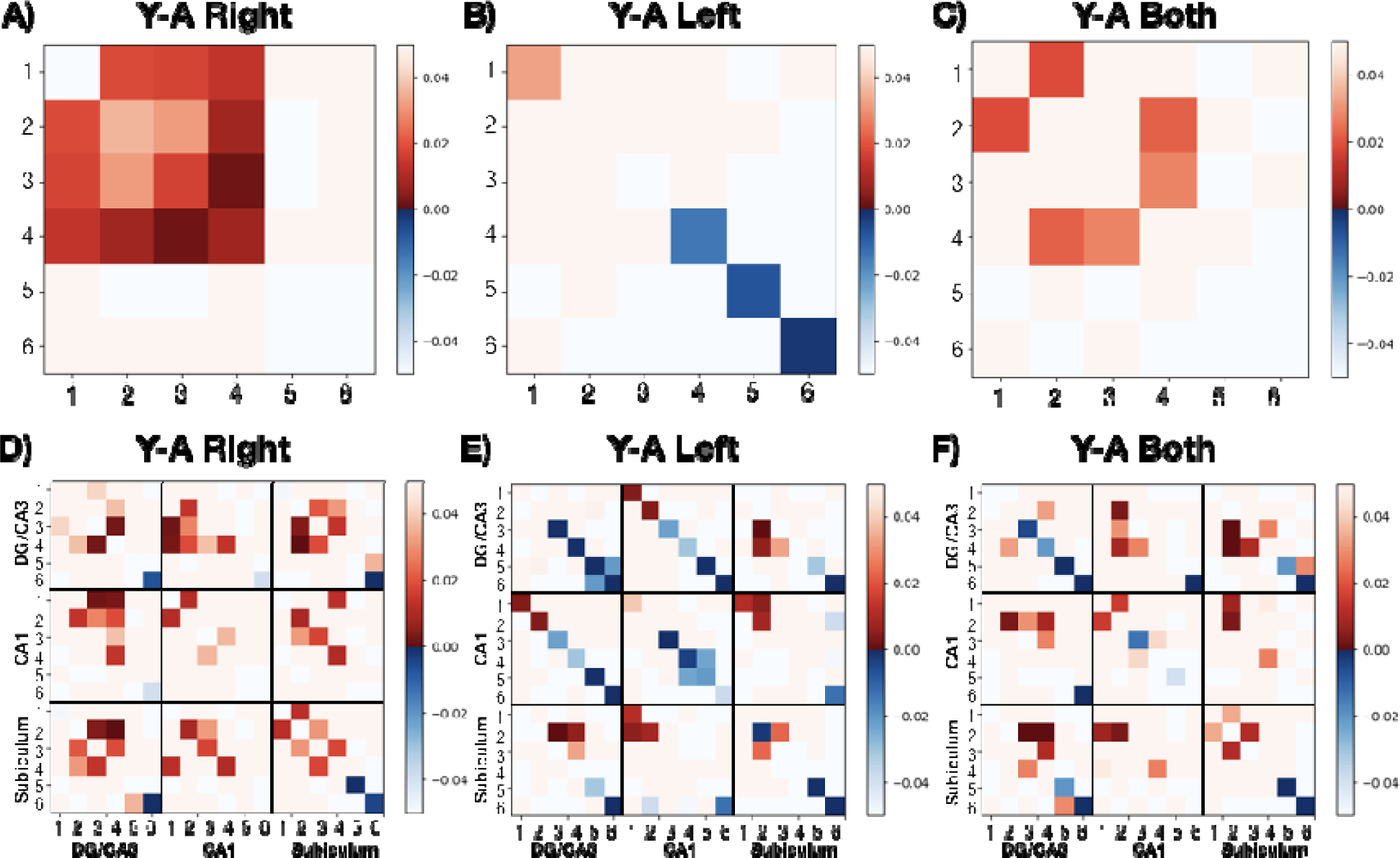
Young-Aged (Y-A) differences in the functional correlation matrices among the 6 segmentations of the hippocampus along the longitudinal axis, reflecting uncorrected p-values with a threshold of p<.05. There is a striking pattern of greater anterior FC in young adults for the whole hippocampus (C), particularly in the right hemisphere (A), and greater posterior FC in older adults, particularly in the left hemisphere (B). When further divided by hippocampal subfield, the same pattern emerges with remarkable regularity in the right (D) and left (E) hemispheres and when averaged together (F).

## 4. Discussion

In this targeted exploration of age-related changes in functional connectivity along the longitudinal axis of the hippocampus to the ERC, PRC, and PHC, we found reliably greater connectivity for younger adults than older ones between the hippocampus and the PHC and, to some degree, in the PRC as well (Figure 2A&B). The drop in PHC FC was more pronounced in the anterior regions of the hippocampus than the posterior ones for older adult and was predictive of lure discrimination performance on the MST, suggesting that this shift may somehow reflect a mechanism that preserves memory performance (Figure 2C&F). Each of the hippocampal subfields exhibited reduced FC anterior regions of the hippocampus in older adults (Figure 4A-C), while the more modest age-related reductions in FC for PRC were more global and not specific to the anterior hippocampus. Finally, we observed reduced inter-hippocampal functional connectivity in older adults in the anterior hippocampus, but greater connectivity in the posterior hippocampus. While age-related dysfunction within the hippocampal subfields has been well-documented, these results suggest that the age-related A-P shift in hippocampal connectivity may also contribute significantly to memory decline in older adults.

### 4.1. A-P hippocampal connectivity: Anterior hippocampal dysfunction

Based on prior reports of age-related changes in anterior hippocampus, altered we had predicted greater anterior than posterior age-related connectivity changes with the PHC. A similar pattern appeared in a brain-wide investigation of FC along the A-P axis of the hippocampus, with reduced connectivity for anterior regions than posterior ones in older adults (48). Indeed, we found reduced connectivity between anterior hippocampus and PHC in older adults (Figure 2B). A-P connectivity within the hippocampus has been shown to be modulated by APOE status, a genetic risk factor for Alzheimer’s disease, with reduced anterior hippocampal connectivity to cortical regions involved in a memory encoding and retrieval task for APOE positive carriers, with no differences in posterior hippocampal connectivity (78). Likewise, across a longitudinal sample, anterior MTL (primarily hippocampal) functional connectivity declined with age, while posterior MTL connectivity remained largely constant (79). There is also evidence for reduced activation in the anterior hippocampus for older adults during an associative recognition memory task (80) and a significant association between volume and memory decline in the anterior hippocampus (81). Thus, reduced anterior hippocampal connectivity to PHC in aging is consistent with declines in anterior hippocampal functioning.

Here, we observed a decrease in FC within the anterior hippocampus in older adults (Figure 5) that is also consistent with an age-related alteration in anterior hippocampal functioning. Unexpectedly, we found some evidence that this may be associated with greater posterior within-hippocampal connectivity (Figure 5). We should note that this pattern could be viewed as either enhanced posterior connectivity in the older sample or as reflecting a de-differentiation of connectivity in the older adults. Regardless, oddly, these competing patterns were asymmetrical, with greater age-related anterior dysfunction in the right hemisphere and greater poster engagement in the left hemisphere. The reason for this discrepancy is not clear, though it is possible that task demands (here, incidental encoding of everyday objects) biased these activity profiles. Again, this age-related anterior hippocampal dysfunction is consistent with reduced volumes in the anterior hippocampus with age reflecting neural deterioration in this region (82).

Interestingly, while there was some evidence for an age-related decline in functional connectivity between the hippocampus and PRC, there was not a difference along the longitudinal axis and the main effect was less reliable. While these findings are consistent with the A-P invariant drop in functional connectivity to default mode network regions in older adults (83), they do not parallel our PHC findings. Before speculating too deeply on this dissociation though, we will review the A-P functional connectivity within the hippocampal subfields.

### 4.2. Age-related dysfunction in A-P functional connectivity for the hippocampal subfields

We had predicted reduced connectivity of DG/CA3 given age-related functional activity dysfunction in this region, which we observed in connectivity to both PHC and PRC. While functional connectivity to PHC was significantly reduced only in the anterior regions for older adults, there was a strong trend for a reduction in posterior hippocampus as well (Figure 4A), suggesting it is hardly so selective. This pattern was mirrored in the functional connectivity to PRC (Figure 4D), offering support for global DG/CA3 dysfunction across the longitudinal axis of the hippocampus. Similarly, functional connectivity between the subiculum (Figure 4C&F) and CA1 (Figure 4B&E) both PHC exhibited anterior-specific declines in older adults, consistent with anterior hippocampal dysfunction noted earlier. Notably, age-related declines in FC with PRC was not anterior-specific for any of the hippocampal subfields.

An earlier investigation of hippocampal subfield connectivity along the longitudinal axis with PRC and PHC in young adults revealed greater anterior FC with PRC and greater posterior FC with PHC, which was relatively consistent across the three hippocampal subfields (35). Here, we found that FC with both PHC and PRC was greater for the anterior hippocampal regions in young adults, approximately following a skewed inverted U-shape function, with generally greater connectivity in the body than either the head or the tail (Figure 2 A&B). This discrepancy, and the lack of age-related anterior specificity in FC to the PRC, may be due, at least in part, to our use of task-based fMRI versus the resting-state fMRI used by Libby et al. (35). While we demonstrated our pattern of results across three datasets, they all involved the encoding of visual objects or scenes, which are known to modulate activity in the hippocampus, PRC, and PHC. Whereas resting-state fMRI suffers from potential pitfalls of being unconstrained, particularly when comparing groups that may engage in different types of rest activities, our task may have biased these functional connectivity relationships due to the active engagement of these regions.

Consistent with prior histological (84) and structural imaging studies (85, 86), we found an uneven distribution of the hippocampal subfields across the longitudinal axis (Figure 4G-I). We should note that there is significant variability in how these regions are segmented across labs, which will induce variance in findings until a validated, standardized protocol to minimize this variability is in place (87). As the subfield representation varies across the long axis, variations in segmentation protocols can influence long-axis findings. Accepting this caveat, we found that age-related volume loss was not consistent across the longitudinal axis of the hippocampus. We found significant volume decline in the posterior regions for both DG/CA3 and the subiculum, but no volumetric decline in anterior regions (but see 82). While the tail of the hippocampus can be notoriously difficult to segment, these volume differences were not limited to the final tail region but begin mid-body of the hippocampus. However, despite these posterior volumetric declines, FC was much weaker in the anterior portions of the hippocampus. This disparity emphasizes two key points: 1) anatomical volume and connectivity does not exclusively dictate hippocampal function and coordination with other regions, and 2) disparities in function across the longitudinal axis of the hippocampus remain consistent despite differences in the inherent properties of the various hippocampal subfields. The differences in hippocampal functional connectivity are not driven by one subfield alone or due to the disparity in subfield volume across the longitudinal axis.

### 4.3. MTL network dysfunction in aging

While there is clear age-related dysfunctiona along the longitudinal axis of the hippocampus and neurobiological changes specific to the hippocampal subfields, there is also substantial evidence for age-related change in the surrounding medial temporal lobe cortices. Ranganath & Ritchey (2) have proposed two cortical systems for memory-guided behavior: an anterior temporal (AT) system that includes the PRC and a posterior medial (PM) system that includes the PHC. There is existing evidence for age-related decline in the AT system, with lower firing rates in aged PRC neurons (88), lower glutamate levels, reflecting reduced excitatory activity in the aged PRC (89), and reduced BOLD activity in the PRC of older adults (90, 91). Aged animals (92) and humans (91) have difficulty discriminating between complex stimuli, consistent with the false recognition of novel stimuli in animals with PRC lesions (93). There is less existing evidence for age-related neurobiological alterations in the PM system and the PHC, but there has been documented volumetric decline (39, 66) and decreases in white matter integrity (94). Here, however, our functional connectivity findings were most robust in this PM system. Together, the literature then indicates some form of age-related dysfunction in both the AT and PM systems (Burke et al., 2018). In sum, our investigation can be summarized thusly:

#### Age-related DG/CA3 alterations dominate network dysfunction

Irrespective of the longitudinal axis, age-related connectivity reductions were most pronounced in DG/CA3 (Figure 3D), both with connectivity to the other hippocampal subfields and to the PHC and PRC. These findings are consistent with models of age-related neurobiological changes in these regions (40) and emphasize the critical nature of these regions for age-related decline (46).

#### Anterior hippocampus shows greater functional connectivity declines

While there is some evidence for posterior hippocampus decline as well, there is clear evidence for anterior hippocampal decline across the subfields to both PHC and PRC and within the hippocampus. These findings are consistent with other reports of anterior hippocampal dysfunction with age, but there is still considerable variability in the literature on this front, due in no small part to differences in functional connectivity regions (default mode network vs other cortical and subcortical regions), resting-state vs task-based fMRI, aging vs APOE-positive or MCI participants, among others. Nevertheless, possible anterior-specific declines are particularly important when considering neuronal recordings and anatomical resections from the hippocampus, which often report from dorsal (posterior) regions since the ventral (anterior) hippocampus can be difficult to access in rodents.

#### Age-related declines in FC from hippocampus to PHC and PRC

Despite clear differences in the functioning of PHC and PRC and the corresponding pathways that they are integrated with (2), both suffer from age-related declines in functional connectivity. These findings are consistent with the model proposed by Burke et al. (2018), which argues for distinctions between these pathways, but showing age-related alterations along both as well. Interestingly, we found age-related impairment in functional connectivity between the hippocampus and these regions, but also between PRC and PHC (Figure 3D). Notably, during the continuous recognition paradigm, the decreased FC between PRC and PHC disappears, suggesting a role for task demands on this relationship (Supplemental Figure 2C). Future studies may probe this relationship in functional connectivity during encoding, retrieval, and non-mnemonic processing.

Despite possible predictions for comparable ERC functional connectivity declines in aging, possibly even dissociating lateral ERC and medial ERC (95, 96), we did not observe anything notable here in any of our datasets. ERC is particularly vulnerable to distortions and signal drop-out in functional MRI due to its proximity to the nasal cavity, which may be exacerbated by our high-resolution imaging protocol. Further, our use of task-based fMRI may not have promoted differential connectivity between the hippocampus and ERC to allow us to detect age-related differences. However, we did find connectivity between ERC and hippocampus for both young and older adults that did not vary across the longitudinal axis of the hippocampus, with an average connectivity within the range of that observed for PHC in older adults. We did explore hippocampal functional connectivity to medial ERC and lateral ERC but did not find any differences in connectivity for younger or older adults, so combined them for improved signal to noise in the analyses presented here.

### 4.4. Concerns and limitations

We exclusively analyzed task-based datasets, reasoning that resting-state functional connectivity suffers from more uncontrolled sources of variability that may or may not be consistent within groups. The possibility that different groups may systematically engage in different rest behaviors is a potential confound for all resting-state FC studies. Even when a cognitive task is regressed out and only the residuals are considered, “state” effects may remain. Here, we only considered task-based functional connectivity but have the following caveat: all of our datasets involved encoding of objects or scenes, which we know can be negatively influenced by age (90, 91, 97). Unfortunately, we do not have access to high-resolution data collected during a non-mnemonic task, but advocate replicating this approach in the future to explore the A-P differentiation of the hippocampus and its subfields. Likewise, our high-resolution scan parameters did not allow for whole-brain imaging, restricting our analyses to the medial temporal lobe network, but newer scan sequences will now allow for whole-brain connectivity analyses at high-resolution to further explore the A-P differentiation of the hippocampus and its subfields.

## 5. Conclusions

We sought to investigate the age-related changes in functional connectivity along the longitudinal axis of the hippocampus to the ERC, PRC, and PHC. We found reliably greater connectivity for younger compared to older adults with the PHC and, to a lesser degree, with the PRC. This drop in functional connectivity was more pronounced in the anterior regions of the hippocampus than the posterior ones for PHC. Interestingly, this A-P shift in older adults was predictive of lure discrimination performance on the MST, suggesting that this shift may reflect a compensation mechanism that preserves memory performance. Each of the hippocampal subfields reflected reduced FC anterior regions of the hippocampus in older adults with some evidence for a posterior decline as well, particularly in DG/CA3. Finally, we observed reduced inter-hippocampal functional connectivity in older adults in the anterior hippocampus, but greater connectivity in the posterior hippocampus. While age-related dysfunction within the hippocampal subfields has been well-documented, these results suggest that the age-related A-P shift in hippocampal connectivity may also contribute significantly to memory decline in older adults.

## Acknowledgements

This research was supported in part by a grant from the National Institutes of Aging: AG034613 and P50 AG016573 (subaward). We thank Samantha Rutledge for assistance in data collection and Branden Kolarik for helpful comments.

## Supplementary Information Text

### Continuous Recognition Task

A total of 23 young (13F/10M; mean age = 27.5; range = 21-34 years) and 25 older adults (14F/11M; mean age = 70.2; range = 59-84 years). Five of these older adults also participated in the primary dataset but viewed a different set of objects during this scan session and it took place approximately 3 years later. Instead of performing the indoor/outdoor task, here participants performed and old, similar, new judgement on repeat, lure, and new items. The scan parameters and data processing pipeline paralleled our primary dataset.

To evaluate whether connectivity along the long axis of the hippocampus differed for young and older adults, we calculated FC along the 6 hippocampal segmentations to three separate medial temporal lobe regions, averaging across left and right hemispheres: ERC, PRC, and PHC. These data were entered into a 2×6 repeated-measures ANOVA with age (young or aging) and region (1,2,3,4,5,6) as variables for each region. All three regions (ERC, PRC, and PHC) showed a main effect across the six hippocampal subdivisions (main effect of region: ERC: F(5,230) = 3.2, p<.01; PRC: F(5,230) = 5.9, p<.0001; PHC: F(5,230) = 8.5, p<.0001). While ERC showed no main effect of age, there was a significant interaction (F(5,230) = 3.2, p<.01). However, Sidak’s multiple comparisons tests on each of the 6 regions did not result in any significant age differences. In constrast, we observed greater connectivity between the Hx and PRC and PHC for younger than older adults (Supplemental Figure 1A-C) (PHC: F(1,46) = 7.0, p <.02). In addition, FC with PHC changed across the six hippocampal subdivisions with an interaction (PRC: F(5,230) = 2.4, p<.05; PHC: F(5, 230) = 2.0, p=.08 (marginal)). Sidak’s multiple comparisons tests on each of the 6 PHC regions revealed greater FC in young than older adults for the first 3 regions (1: t(276) = 3.1, p<.02; 2: t(276) = 2.6, p<.05; 3: (t(276) = 2.7, p <.05), making up the anterior portion of the hippocampus, but not the last 3 regions (4: t(276) = 2.0, p=.23; 5: t(276) = .64, p=.99; 6: t(276) = 1.0, p=.89). While Hx-PRC showed an interaction, none of the six subdivisions showed a reliable effect of age, consistent with the lack of a main effect of age.

### Scene Encoding Task

A total of 34 young (21F/13M; mean age = 28.3; range = 20-39 years) and 33 older adults (19F/14M; mean age = 76.2; range = 70-87 years). Twenty-seven young and 26 older adults also participated in the primary dataset within approximately 6 months as part of a larger study on the neural basis of age-related memory decline. Participants viewed pictures of scenes and determined if each one was oriented in a portrait or landscape orientation while in the scanner. They were later tested on their memory for these images, but these imaging data are from this encoding portion of the task only. The scan parameters and data processing pipeline paralleled our primary dataset.

Again, these data were entered into a 2×6 repeated-measures ANOVA with age (young or aging) and region (1,2,3,4,5,6) as variables for each region. All three regions (ERC, PRC, and PHC) showed a main effect across the six hippocampal subdivisions (main effect of region: ERC: F(5,325) = 3.7, p<.01; PRC: F(5,325) = 7.4, p<.0001; PHC: F(5,325) = 8.4, p<.0001). While ERC and PRC showed no main effect of age or interaction, we observed greater connectivity between the Hx and PHC for younger than older adults (Supplemental Figure 1E-F) (PHC: F(1,65) = 6.7, p <.02). In addition, FC with PHC changed across the six hippocampal subdivisions with an interaction (PHC: F(5, 325) = 2.7, p<.05). Sidak’s multiple comparisons tests on each of the 6 PHC regions revealed greater functional connectivity in young than older adults for 2 of the first 3 regions (1: t(390) = 1.4, p=.69; 2: t(390) = 3.6, p<.01; 3: (t(390) = 2.9, p <.05), making up the anterior portion of the hippocampus, but not the last 3 regions (4: t(390) = .89, p=.94; 5: t(390) = 1.7, p=.43; 6: t(390) = .03, p=.99), consistent with the other datasets demonstrating an age-related decrease in FC to the PHC in the anterior hippocampus relative to the posterior hippocampus.

### Hemispheric Asymmetry in Object Encoding Task

We explored whether there was hemispheric asymmetry in these findings by examining each FC relationship within hemisphere: left Hx to left PHC, PRC, ERC and right Hx to right PHC, PRC, ERC. Again, FC changed across the 6 hippocampal regions for both left and right hemispheres for all MTL cortices (main effect of region: all p’s<.05). The Hx-PHC showed an age related shift in both hemispheres (main effect of age: left: F(1,60) = 8.7, p<.005; right: F(1,60) = 9.4, p<.005) and the Hx-PRC showed a marginal age-related shift only in the right hemisphere (F(1,60) = 3.4, p = .07) with greater FC for young adults than older adults. Again, we collapsed activity into anterior and posterior regions and found reduced Hx-PHC FC in the anterior hippocampus in older adults (right: t(120) = 3.8, p<.001; left: t(120) = 3.3, p<.01) than in the posterior hippocampus (right: t(120) = 2.0, p = .10; left: t(120) = 1.9, p = .11), supported by a significant interaction in the right hemisphere (F(5,300) = 6.8, p<.02). Therefore, there is some evidence for hemispheric asymmetry, with larger effects present in the right than left hemisphere, but both hemispheres consistent in showing an age-related decrease in Hx-PHC functional connectivity.

**Fig. S1.**
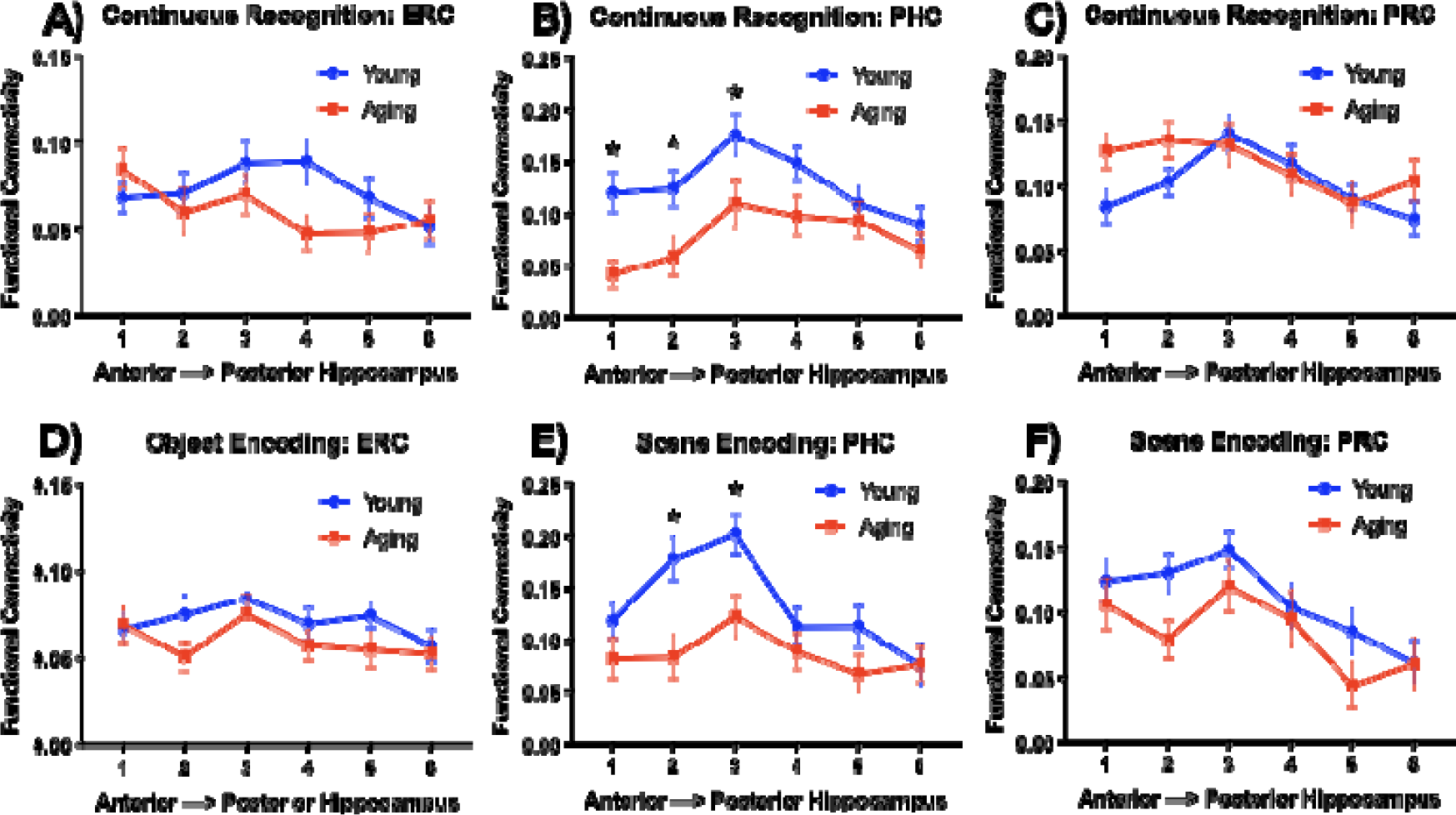
There was no effect of age on functional connectivity with the ERC in any of the datasets, including the continuous recognition task (A) and object encoding task (D). However, functional connectivity across the longitudinal axis of the hippocampus with the PHC shows an age-related decline in the anterior portions during both a continuous recognition task (B) and an incidental scene encoding task (E). There was an overall age-related decline in FC for Hx-PRC during the incidental scene encoding task (F) that was not observed during the continuous recognition task (C), suggesting that task demands may be able to modulate this effect.

**Fig. S2.**
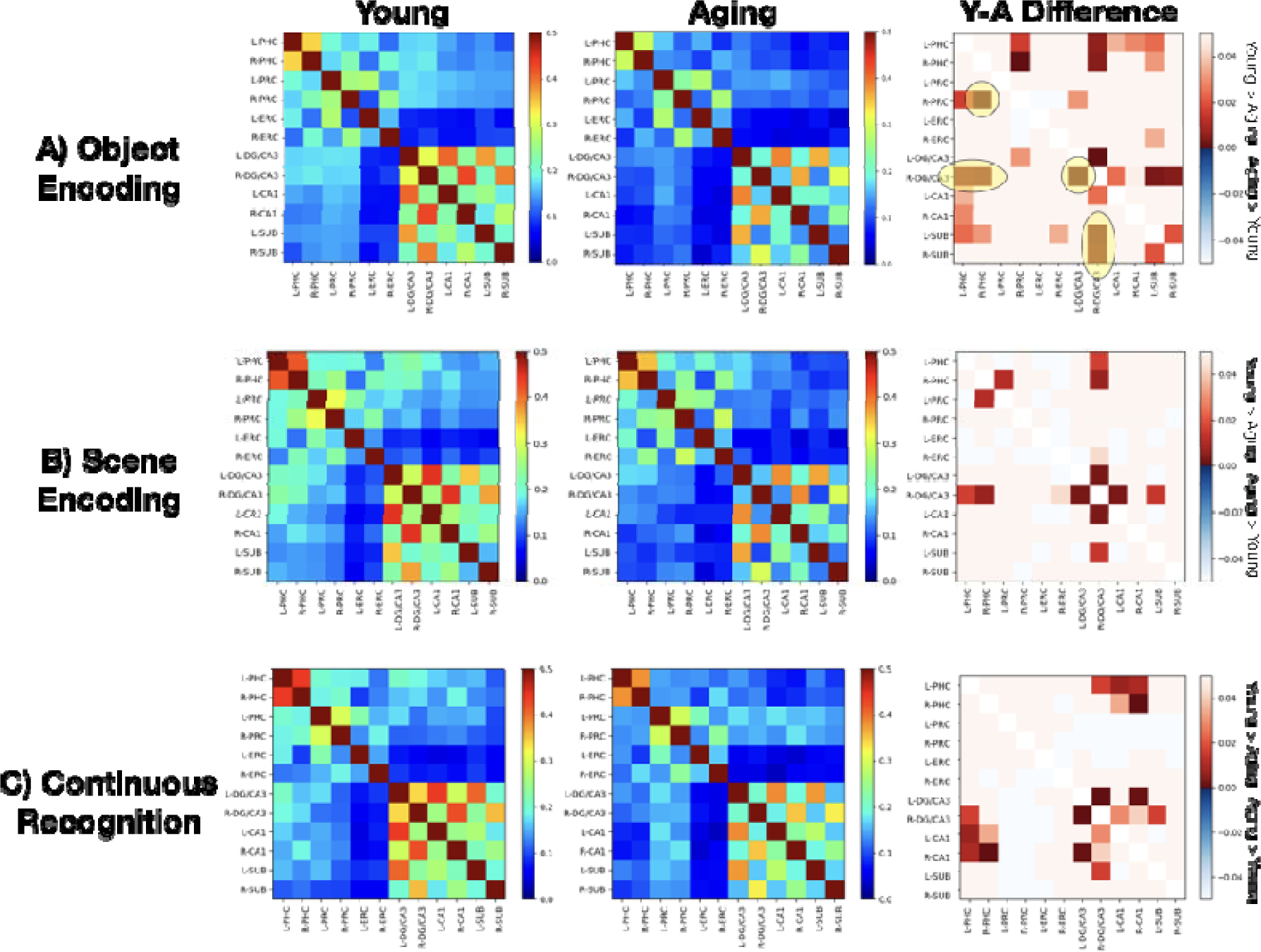
Functional connectivity matrices for young and aging adults and the difference matrix mapping p-values for each of the 3 datasets (object encoding data is identical to Figure 4 but shown again here for direct comparison). Difference matrix testing whether regions showed greater connectivity in young than old or vice-versa. The regions that passed multiple comparisons thresholding in the Object Encoding dataset are circled in yellow, but we did not impose that strict comparison on the other two datasets since we were looking for replication of those a priori regions.

**Fig. S3.**
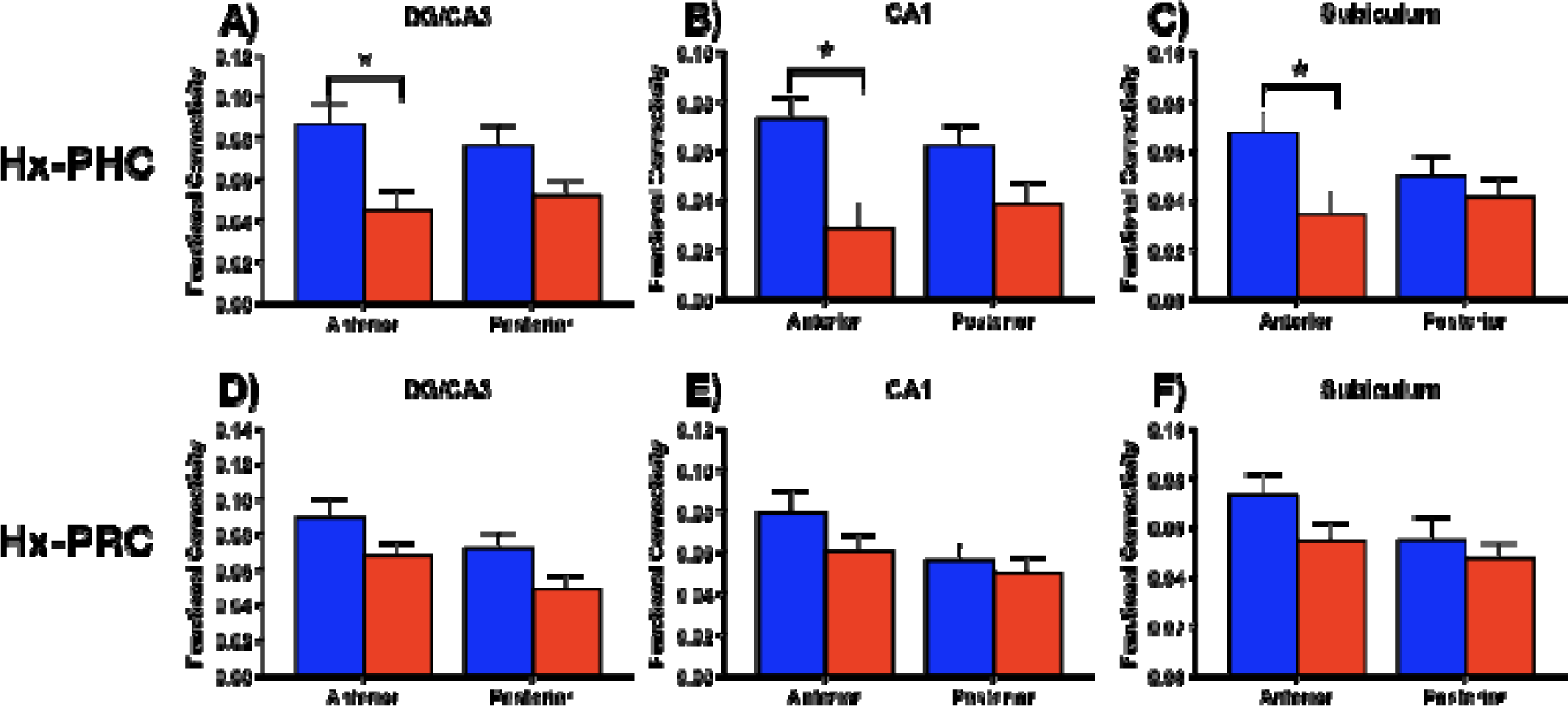
Functional connectivity between Hx-PHC and Hx-PRC was averaged across the first 3 anterior regions and last 3 posterior regions for each of the hippocampal subfields. Both DG/CA3 (A) and Subiculum (C) showed an age-related decline in the anterior region, while CA1 also showed a decline in the posterior region (B). Similarly, CA1 (E) and Subiculum showed an age-related decline in anterior connectivity to PRC, while D

